# A universal stress protein is essential for the survival of *Mycobacterium tuberculosis*

**DOI:** 10.1101/2023.01.23.525157

**Authors:** Arka Banerjee, Moubani Chakraborty, Suruchi Sharma, Ruchi Chaturvedi, Avipsa Bose, Priyanka Biswas, Amit Singh, Sandhya S. Visweswariah

## Abstract

*Mycobacterium tuberculosis* employs several signaling pathways to regulate its cellular physiology and survival within the host. Mycobacterial genomes encode multiple adenylyl cyclases and cAMP effector proteins, underscoring the diverse ways in which these bacteria utilize cAMP. We have earlier identified universal stress proteins (USP), Rv1636 and MSMEG_3811 in *M. tuberculosis* and *M. smegmatis* respectively, as abundantly expressed, novel cAMP-binding proteins. In this study, we show that these USPs may function to regulate cAMP signaling by direct sequestration of the second messenger. In slow-growing mycobacteria, concentrations of Rv1636 were equivalent to the amounts of cAMP present in the cell, and overexpression of Rv1636 in *M. smegmatis* increased levels of ‘bound’ cAMP. Rv1636 is secreted via the SecA2 secretion system in *M. tuberculosis* but is not directly responsible for the efflux of cAMP from the cell. While *msmeg_3811* could be readily deleted from the genome of *M. smegmatis*, we find that the *rv1636* gene is essential for growth of *M. tuberculosis*, and this functionality depends on the cAMP-binding ability of Rv1636. This is the first evidence of a ‘sink’ for any second messenger in bacterial signaling that would allow mycobacterial cells to regulate the available intracellular ‘free’ pool of cAMP.

## INTRODUCTION

The genus *Mycobacterium* harbours a number of diverse species, ranging from free-living members to several pathogenic species, including *M. tuberculosis*, the causative agent of human tuberculosis. Tuberculosis alone was responsible for the death of ~1.6 million people in 2021 (1) It is therefore imperative to understand the physiology of *M. tuberculosis* to identify effective new anti-mycobacterial strategies.

It has now been well-established that cAMP levels are high in mycobacteria (2, 3), and cAMP is also secreted to the extracellular milieu (2–4). As seen in many other pathogenic bacteria, *M. tuberculosis* utilises cyclic AMP-mediated signaling to subvert the host (3, 5), and a remarkably high number of adenylyl cyclases as well as cAMP effector proteins are encoded in its genome (3, 6–8). However, fast-growing non-pathogenic members, such as *M. smegmatis*, also harbour multiple adenylyl cyclases and cAMP-binding proteins (8, 9). Our earlier work has shown that a significant fraction of intracellular cAMP remains protein-bound in both fast and slow-growing mycobacteria (10). Subsequently, we identified specific universal stress proteins (USP), Rv1636 and MSMEG_3811 in *M. tuberculosis* and *M. smegmatis* respectively, as abundantly expressed cAMP-binding proteins in these organisms (10). Cyclic AMP is known to bind to cyclic nucleotide-binding (CNB) or GAF domains (11–15) in proteins. However, identification of Rv1636/MSMEG_3811 added the USP domain to the list of cyclic nucleotide-binding proteins. USPs are found in bacteria, Archaea, fungi, plants and a few invertebrates, and a subset of them are known to bind ATP (16–18). Rv1636/MSMEG_3811 formed a new class of USPs which bound cAMP with higher affinity than ATP (10). Interestingly, Rv1636-like cAMP-binding USPs are restricted to *Mycobacterium, Corynebacterium* and *Nocardia* genera.

Rv1636 and MSMEG_3811 show similar biochemical properties, and both bind cAMP with an affinity of 3 μM with a stoichiometry of 1:1 (10). The crystal structure of cAMP bound MSMEG_3811 revealed how the ATP-binding pocket in the USP-fold can preferentially accommodate cAMP (10). In the present study, we show that Rv1636 is secreted from the cell by the SecA2 pathway, as well as provide evidence that Rv1636/MSMEG_3811 regulate the available pool of cAMP in the mycobacterial cell. Importantly, the essentiality of *rv1636* in *M. tuberculosis* is dependent on the ability of this USP to bind cAMP.

## MATERIALS AND METHODS

### Strains and culture conditions

*Mycobacterium smegmatis* mc^2^ 155 cells were grown in Middlebrook 7H9 medium (BD Biosciences) supplemented with 0.2 % glycerol and 0.05 % Tween 80 at 37 ^o^C with shaking at 200 rpm. *M. bovis* BCG and *M. tuberculosis* H37Rv cultures were grown in the same medium containing OADC (oleic acid-albumin-dextrose-catalase) supplement at a final concentration of 10 % (v/v) in tissue culture flasks at 37 ^o^C in a humidified incubator in biosafety level 2 (BSL2) and BSL3 facilities, respectively. For solid media, *M. smegmatis* cells were grown in Middlebrook 7H10 agar (BD Biosciences) containing 0.5 % glycerol, and *M. tuberculosis* cells were grown in Middlebrook 7H10 agar supplemented with 0.5 % glycerol and 10 % OADC. As and when required, kanamycin and hygromycin were used at 20 and 50 μg/mL concentrations, respectively.

For generation of the SecA2 complemented strain, PCR was carried out using genomic DNA of *M. tuberculosis* H37Rv. The *secA2* gene was amplified in two parts, and PCR amplicons ligated into cloning vector pCR2.1 (Thermo Fisher Scientific, USA). Clones were digested with *Bam*HI and *Kpn*I (5’ region of *secA2)* or *Kpn*I and *Hind*III (3’ send of *secA2)* and cloned into *Bam*H1-*Hind*II digested pMV10-25 vector The pMV10-25 SecA2 complement clone was electroporated into the electrocompetent Δ*secA2* strain and colonies selected on plates containing hygromycin. Colonies were screened by PCR to confirm the complementation of the *SecA2* gene.

A list of all strains used in this study is provided in Table 1.

**Table 1.** List of strains used in this study

| Strain name | Description | References |
| --- | --- | --- |
| *M. smegmatis* | *M. smegmatis* mc<sup>2</sup> 155 electroporation proficient wildtype strain |  |
| *M. smegmatis* VC | *M. smegmatis* wildtype harbouring episomal pMV10-25 plasmid (Hyg<sup>+</sup>) | This study |
| *M. smematis* Rv1636 OE | *M. smegmatis* wildtype harbouring episomal pMV10-25-P<sub>rv1636</sub>-rv1636 plasmid (Hyg<sup>+</sup>) | This study |
| *Δmsmeg_3811* | *M. smegmatis* strain deleted for the endogenous *msmeg_3811* gene (unmarked) | This study |
| *M. tuberculosis* | *M. tuberculosis* H37Rv wildtype strain |  |
| *Mtb::rv1636* | *M. tuberculosis* wildtype strain having an integrated copy of *rv1636* gene at the L5 *att* site as pMV306-P<sub>rv1636</sub>-rv1636 (Kan<sup>+</sup>) | This study |
| *Mtb::rv1636<sup>DM</sup>* | *M. tuberculosis* wildtype strain having an integrated copy of *rv1636<sup>DM</sup>* allele at the L5 *att* site as pMV306-P<sub>rv1636</sub>-rv1636<sup>DM</sup> (Kan<sup>+</sup>) | This study |
| *Δrv1636::rv1636* | *M. tuberculosis* strain deleted for the endogenous *rv1636* gene, but harbouring an additional copy at the L5 *att* site as pMV306-P<sub>rv1636</sub>-rv1636 (Kan<sup>+</sup>) | This study |
| *ΔRD1* | *M. tuberculosis* H37Rv strain deleted for the *RD1* locus | (62, 63) |
| *ΔsecA2* | *M. tuberculosis* H37Rv strain deleted for the *secA2* gene | (41) |
| *Mtb::cfp10-nluc* | *M. tuberculosis* wildtype strain harbouring integrated copy of pMV306-P<sub>cfp10</sub>-cfp10-nluc at the L5 *att* site (Kan<sup>+</sup>) | This study |
| *ΔsecA2::secA2* | *M. tuberculosis* H37Rv *ΔsecA2* strain complemented with the wildtype *secA2* gene | This study |
| *Mtb::rv1636-nluc* | *M. tuberculosis* wildtype strain harbouring integrated copy of pMV306-P<sub>rv1636</sub>-rv1636-nluc at the L5 *att* site (Kan<sup>+</sup>) | This study |
| *Mtb::nluc* | *M. tuberculosis* wildtype strain harbouring integrated copy of pMV306-P<sub>rv1636</sub>-nluc at the L5 *att* site (Kan<sup>+</sup>) | This study |
| ***ΔRD1::cfp10-nluc*** | *M. tuberculosis ΔRD1* strain harbouring integrated copy of pMV306-P<sub>cfp10</sub>-*cfp10-nluc* at the L5 att site (Kan<sup>+</sup>) | This study |
| ***ΔRD1::rv1636-nluc*** | *M. tuberculosis ΔRD1* strain harbouring integrated copy of pMV306-P<sub>rv1636</sub>-*rv1636-nluc* at the L5 att site (Kan<sup>+</sup>) | This study |
| ***ΔRD1::nluc*** | *M. tuberculosis ΔRD1* strain harbouring integrated copy of pMV306-P<sub>rv1636</sub>-*nluc* at the L5 att site (Kan<sup>+</sup>) | This study |
| ***ΔsecA2::cfp10-nluc*** | *M. tuberculosis ΔsecA2* strain harbouring integrated copy of pMV306-P<sub>cfp10</sub>-*cfp10-nluc* at the L5 att site (Kan<sup>+</sup>) | This study |
| ***ΔsecA2::rv1636-nluc*** | *M. tuberculosis ΔsecA2* strain harbouring integrated copy of pMV306-P<sub>rv1636</sub>-*rv1636-nluc* at the L5 att site (Kan<sup>+</sup>) | This study |
| ***ΔsecA2::nluc*** | *M. tuberculosis ΔsecA2* strain harbouring integrated copy of pMV306-P<sub>rv1636</sub>-*nluc* at the L5 att site (Kan<sup>+</sup>) | This study |
| ***Mtb::dCas9*** | *M. tuberculosis* wildtype strain harbouring integrated copy of pRH2502 plasmid encoding *S. pyogenes* dCas9 (Kan<sup>+</sup>) | (27) and this study |
| ***Mtb::dCas9/scrambled-sgRNA*** | *Mtb::dCas9* strain harbouring the pRH2521 plasmid (Kan<sup>+</sup> Hyg<sup>+</sup>) | (27) and this study |
| ***Mtb::dCas9/rv1636-sgRNA*** | *Mtb::dCas9* strain harbouring the pRH2521-rv1636+11 plasmid (Kan<sup>+</sup> Hyg<sup>+</sup>) | This study |

For experiments to observe the effect of InterBioScreen (https://www.ibscreen.com) natural compounds on *M. tuberculosis* and *M. smegmatis*, cultures were grown to log phase in nutrient-rich media and 5 μl of the culture (or dilutions as indicated) were spotted on 7H9 agar media containing 10 % OADC enrichment, 0.2 % glycerol, with or without the appropriate compound at the indicated concentrations. Plates were photographed after 7-10 days of incubation for *M. tuberculosis* and 2-3 days for *M. smegmatis*.

### Immunoblotting

Protein samples were electrophoresed on SDS-polyacrylamide gel of appropriate percentage and transferred to PVDF membrane for immunoblotting. Blots were probed with primary antibody in TBST (10 mM Tris-Cl (pH 7.5), 0.9 % NaCl and 0.1 % Tween 20) containing 0.2 % BSA (TBT) overnight at 4 °C followed by horseradish peroxidase (HRP)-conjugated secondary anti-rabbit IgG antibody (1:50,000) for 1 h at room temperature. Bound antibody was detected using enhanced chemiluminescence with Immobilon Western HRP Substrate (Merck Millipore, USA). 20-25 μg of protein was loaded for whole cell lysates and sub-cellular fractions, whereas 25 ng of protein was loaded for purified recombinant Rv1636. Antisera against Rv1636 was raised by immunising a rabbit with purified, recombinant Rv1636. Anti-CRP (19), anti-CFP-10 (from BEI Resources, NIAID, National Institutes of Health, USA) were used at recommended dilutions. Anti-CRP and anti-GyrB antibodies were described earlier and available in the laboratory (19, 20).

### Preparation of culture filtrate, cell envelope and cytosolic fractions

For preparation of culture filtrate from different strains of *M. tuberculosis* H37Rv cultures, cells were grown in 100 mL of 7H9 medium containing 0.2 % glycerol in roller culture bottles for 4 weeks at 37 °C. Cells were harvested by centrifugation, culture supernatant was collected and filtered through a 0.2 μm syringe filter. Culture supernatant proteins were precipitated using 20 % trichloroacetic acid (TCA) at −20 °C overnight, and centrifuged at 20,000 x *g* at 4 °C for 1 h. Proteins were then washed with ice cold acetone twice and pellets air-dried and resuspended in 150 mM Tris-Cl (pH 8) (21). Equal volumes of the resuspended protein for different strains were mixed with SDS loading dye, heated at 95 °C for 10 minutes, and loaded for analysis by SDS-PAGE followed by Western blotting.

The cell pellets were washed with cold phosphate-buffered saline (PBS) and resuspended in lysis buffer containing 50 mM Tris-Cl (pH 8.2), 100 mM NaCl, 10 mM 2-mercaptoethanol (2-ME), 10 % glycerol, and 1 mM phenylmethylsulphonyl fluoride (PMSF). Cells were lysed by bead beating, centrifuged at 1000 x *g* for 1 minute, followed by harvesting of the supernatant and further centrifugation at 17,000 x *g* for 20 minutes at 4 °C. The supernatant fraction was used as the cytosol, while the pellet was resuspended in fresh lysis buffer and used as the cell envelope fraction. Protein samples were quantified by Bradford method and equal protein (20-25 μg) was loaded for SDS-PAGE followed by immunoblotting.

### Monitoring secretion of Rv1636 using a nanoluciferase assay

Nanoluciferase (Nluc) gene was obtained from the pNL1.1 plasmid (Promega Corporation, USA). The nanoluciferase encoding gene was fused to the C-terminus of either CFP-10 or Rv1636 following a short linker sequence (GGSGGGSSGG) as described earlier for other fusion constructs (22). The *cfp10-nluc* and *rv1636-nluc* constructs were cloned under the respective endogenous *cfp-10* and *rv1636* promoters in the pMV306 vector. As a control, the *nluc* gene alone was cloned under the *rv1636* promoter. Each construct was individually electroporated in wildtype, *ΔRD1* and *ΔsecA2 M. tuberculosis* strains (Table 1). The strains were grown in 7H9 medium containing 0.2 % glycerol, 0.05 % Tween 80 and 10 % OADC supplement with 20 μg/mL kanamycin till OD_595nm_ reached ~1 in 24-well plates. The cells were pelleted, washed twice with fresh medium and finally resuspended in fresh 7H9 medium containing 0.2 % glycerol, 0.05 % Tween 80 and 20 μg/mL kanamycin such that the OD_595nm_ was 1. The cultures were incubated at 37 °C, harvested at different times and culture supernatants collected by centrifugation of the suspension at 12,000 x *g* for 5 minutes at room temperature.

For measuring Nluc activity, culture supernatants were assayed using 2.5 μM coelenterazine 400a as substrate. The assay buffer was essentially used as described by Hall *et al*. and consisted of 50 mM MES (pH 6), 1 mM EDTA, 0.5 % NP-40, 150 mM KCl, 1 mM DTT, 35 mM thiourea, 2.5 μM coelenterazine 400a (22). For the assay, 40 μL of culture supernatant was mixed with equal volume of 2X assay buffer containing coelenterazine 400a, incubated for 3 min at RT, and luminescence counts were measured in a Tecan Infinite (m200 PRO) plate reader. Luminescence counts at 0 h were subtracted from other readings, and secretion of Nluc was represented as the fold increase in luciferase activity of Rv1636-Nluc fusion strains to that of the Nluc control.

### Cyclic AMP measurement

Cyclic AMP measurements were performed using radioimmunoassay as described previously (2, 10, 23, 24). Briefly, for separation of intracellular bound and free cAMP, cells were harvested as the cells started entering stationary phase (OD_595nm_ ~2-3). The cell pellets were washed with cold PBS, resuspended in lysis buffer containing 50 mM Tris-Cl (pH 8.2), 100 mM NaCl, 10 mM 2-ME, 10 % glycerol, and 1 mM PMSF, and lysed by bead beating. The lysates were centrifuged at 1000 x *g* for 1 minute, followed by harvesting of the supernatant and further centrifugation at 17,000 x *g* for 10 minutes at 4 °C. Next, protein (400 μg) in the supernatant (cytosolic fraction) was subjected to centrifugation through a 3 kDa cut-off membrane filter (Amicon Ultra-0.5 3-kDa Ultracell, Millipore) at 4 °C. Cyclic AMP was measured from the eluate (“free” cAMP) and neat cytosolic fractions (“total” cAMP), while the “bound” cAMP fraction was estimated by subtracting the “free” cAMP from the “total” cytosolic cAMP (10, 23).

Culture supernatant was used to measure the total extracellular cAMP concentrations. Cyclic AMP was measured by radioimmunoassay following acidification and acetylation of all samples.

### ATP measurement

ATP contents were estimated using the ATP Determination Kit (Catalogue number A22066; Thermo Fisher Scientific, USA). For preparation of samples, 200 μL of culture was collected and cells pelleted. Cells were resuspended in an equal volume of deionized water, and boiled at 95 °C for 10 minutes. Samples were stored at −70 °C until measurement. 8 μL samples were used in an 80 μL reaction to determine ATP levels against an ATP standard curve following the manufacturer’s protocol. Luminescence was measured in a white 96-well plate using Tecan Infinite (m200 PRO) plate reader. ATP measurements were normalised for protein content in the samples as determined by Bradford method.

### Southern blotting

Genomic DNA of *M. tuberculosis* strains were prepared as described earlier (25). Southern blotting to confirm the genotypes were performed using ECL™ Direct Nucleic Acid Labeling and Detection kit (GE Healthcare, USA). Probe (650 bp) specific to the *rv1636* allele (Fig 6A) was prepared by PCR using Rv1636 5’KO Fwd and Rv1636 promoter Rev primers, and non-radioactively labelled with horseradish peroxidase (HRP) according to manufacturer’s protocol. Hybridization was carried out at 42 °C post capillary transfer of *Kpn*I digested *M. tuberculosis* gDNA to Hybond-N+ nylon membrane (GE Healthcare, USA). The blot was developed using chemiluminescent substrates (Immobilon Western HRP Substrate, Merck Millipore, USA). Sequences of all primers used in this study are shown in Table 2.

**Table 2.**
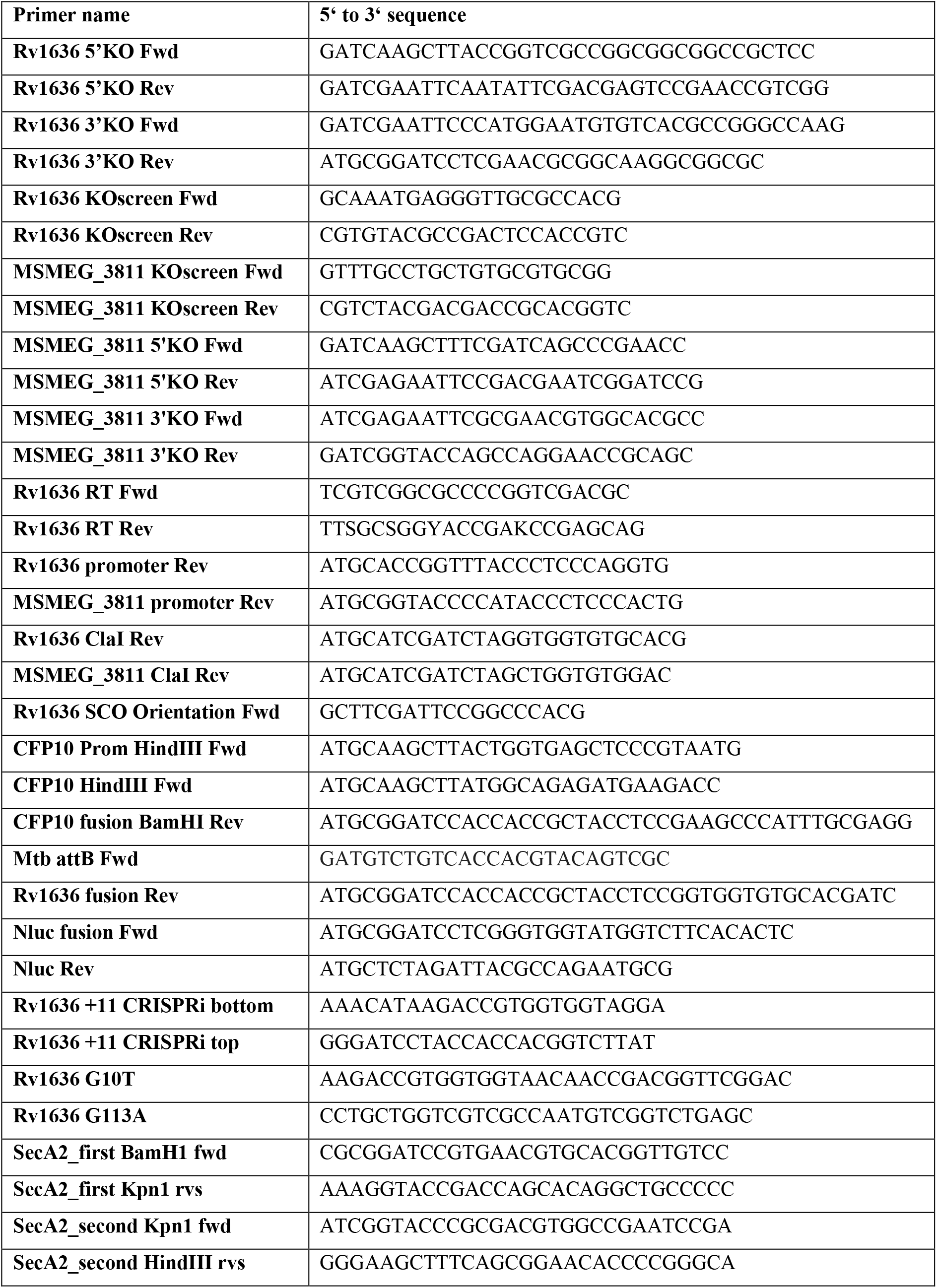
List of primers used in this study

### Generation of *Δmsmeg_3811* and *Δrv1636* strains

The *msmeg_3811* gene was deleted in wildtype *M. smegmatis* strain following the protocol as described by Parish and Stoker (26). For generation of the knockout, 650 nucleotides upstream and downstream of *msmeg_3811* and 51 nucleotides from the start and before the stop codon of the *msmeg_3811* ORF, respectively, were cloned into the p2NIL vector to generate the p2NIL-*msmeg_3811*KO construct. The *Pac*I fragment from pGOAL19 containing three markers (β-galactosidase, hygromycin resistance, and sucrose susceptibility (*sacB*)) was cloned into p2NIL-*msmeg_3811*KO to generate plasmid p2NIL-*msmeg_3811*KO-*Pac*I. 5-10 μg of p2NIL-*msmeg_3811*KO-*Pac*I was electroporated into wildtype *M. smegmatis* electrocompetent cells. Single crossover (SCO) colonies were selected using kanamycin, hygromycin and X-gal markers. Single crossovers were allowed for the second crossover and the positive colonies were selected using sucrose and X-gal. The double crossovers were further confirmed using genomic PCR and immunoblotting (Fig 4B).

For deletion of the *rv1636* gene from wildtype *M. tuberculosis* H37Rv, a similar two-step selection strategy was used as that for *msmeg_3811* knockout generation (26). 650 nucleotides upstream and downstream of *rv1636* and 51 nucleotides from the start and before the end of the *rv1636* ORF, were cloned into the p2NIL vector to generate p2NIL-*rv1636*KO construct. Two merodiploid strains were generated by integrating either a wildtype copy of *rv1636* gene or an *rv1636^DM^* allele at the L5 *att* site of the *M. tuberculosis* genome, to generate *Mtb::rv1636* and *Mtb::rv1636^DM^* strains, respectively. Single crossover (SCO) colonies were confirmed by PCR and Southern blotting (Fig 6B), and positive colonies were proceeded with for the second crossover. All double crossover (DCO) colonies appearing at the end of the process were screened by PCR using specific primers for the deletion of the endogenous *rv1636* gene (Fig 6A).

### CRISPRi-mediated knockdown of *rv1636*

CRISPRi-based knockdown of *rv1636* in *M. tuberculosis* H37Rv was performed using the strategy as originally described by Singh *et al*. (27). First, the codon-optimised *S. pyogenes* dCas9 expressing integrative plasmid pRH2502 was introduced into wildtype *M. tuberculosis* H37Rv to generate the strain *Mtb::dCas9*. A 20 nt-long *rv1636* targeting spacer sgRNA sequence (+11 to +30 position within the *rv1636* ORF) was cloned within the two *Bbs*I sites in the pRH2521 plasmid to generate pRH2521-Rv1636+11 construct. Either the episomal pRH2521 (scrambled non-targeting control) or pRH2521-Rv1636+11 plasmids were transformed into *Mtb::dCas9* strain to generate *Mtb::dCas9/scrambled-sgRNA* and *Mtb::dCas9/rv1636-sgRNA* strains, respectively. Cultures were grown in Middlebrook 7H9 broth, supplemented with 0.2 % glycerol, 10 % OADC (oleic acid, albumin, dextrose and catalase), 0.05 % Tween, 50 μg/ml hygromycin and 25 μg/ml kanamycin, under static conditions at 37^0^C. Anhydrotetracycline (ATc, 100 ng/mL) was added to the media and replenished every 48 h. To assess the viability of cultures, 5 μl of culture was spotted on 7H10 agar media containing 10 % OADC enrichment and 0.2 % glycerol and incubated for 6-7 days.

Suspension cultures were harvested 2 and 4 days after treatment with ATc, lysates prepared and subjected to western blot analysis with an antibody to Rv1636 as described above.

### Biolayer interferometry

Protein binding to cAMP monitored using biolayer interferometry (BLI) was carried out on the Octet RED96 system (Pall ForteBio LLC, USA). Recombinant Rv1636^DM^ (Rv1636^G10T G113A^) was generated by site-directed mutagenesis taking pPRO-Rv1636 plasmid as the template (10, 28). Both recombinant Rv1636 and Rv1636^DM^ proteins were expressed and purified from *E. coli* SP850 *cyc*^-^ strain (29) as described previously (10). For BLI, 1 mM 8- (6-Aminohexylamino) cAMP (8-AHA-cAMP; BIOLOG) prepared in 100 mM HEPES (pH 8) was used for conjugation (1200 s) to AR2G tips (amine reactive) after activation (720 s) with a mixture of 20 mM EDC and 10 mM NHS in a final reaction volume of 200 μL. The reaction was quenched by addition of 1 M ethanolamine-HCl (pH 8.5). Reference tips were activated and deactivated without the addition of 8-AHA-cAMP. All interactions were studied in buffer containing 10 mM HEPES (pH 7.5), 100 mM NaCl, 10 mM 2-ME, 10 % glycerol, 0.5 % BSA and 0.05 % Triton X-100. All proteins were used at a concentration of 4 μM. The association and dissociation were both monitored for 10 minutes at 25 °C. After interaction, the tips were regenerated with 0.05 % SDS prepared in the assay buffer. Binding kinetics were analysed using Octet Data Analysis software, version 8.2 (Pall ForteBio LLC, USA).

### Statistical analyses

All graphs were plotted and statistical analyses were performed using GraphPad Prism 7.

## RESULTS

### Genomic organisation of *msmeg_3811/rv1636*

Our earlier report identifying MSMEG_3811 and Rv1636 in *M. smegmatis* and *M. tuberculosis*, respectively, as cAMP-binding proteins (10) added to the repertoire of cAMP-binding proteins present in mycobacteria. Additionally, the fact that MSMEG_3811 and Rv1636 bind cAMP through the USP-fold (10) warranted further investigation into the cellular functions of these universal stress proteins (USP) in mycobacterial physiology. First, we studied the genomic locus encoding *rv1636* and its orthologs across different mycobacterial species (Fig 1A). In all members of the *M. tuberculosis* complex (MTBC), *rv1636* and its orthologs are encoded as a monocistron, with identical organisation of flanking genes in this locus (Fig 1A). Interestingly, in *M. leprae*, the *rv1636* ortholog, *ml1390*, is predicted to be a functional gene with conservation of the genomic locus (Fig 1A). In non-tuberculous species such as *M. marinum* and *M. avium*, the orthologous genes are also encoded as a monocistron (Fig 1A). The *msmeg_3811* gene in fast-growing *M. smegmatis* is likely to be expressed as a monocistron, while a distinct operon encoding two genes, *msmeg_3813* and *msmeg_3812*, is located upstream of *msmeg_3811* as predicted by the database of prokaryotic operons (DOOR) (30) (Fig 1A). Further, in all mycobacterial species, the genes are orientated in the direction of the movement of the replication fork. This overall similar genomic organisation of *rv1636* and its orthologs suggests similar modes of regulation, expression and perhaps function of this protein across different mycobacterial species.

**Figure 1.**
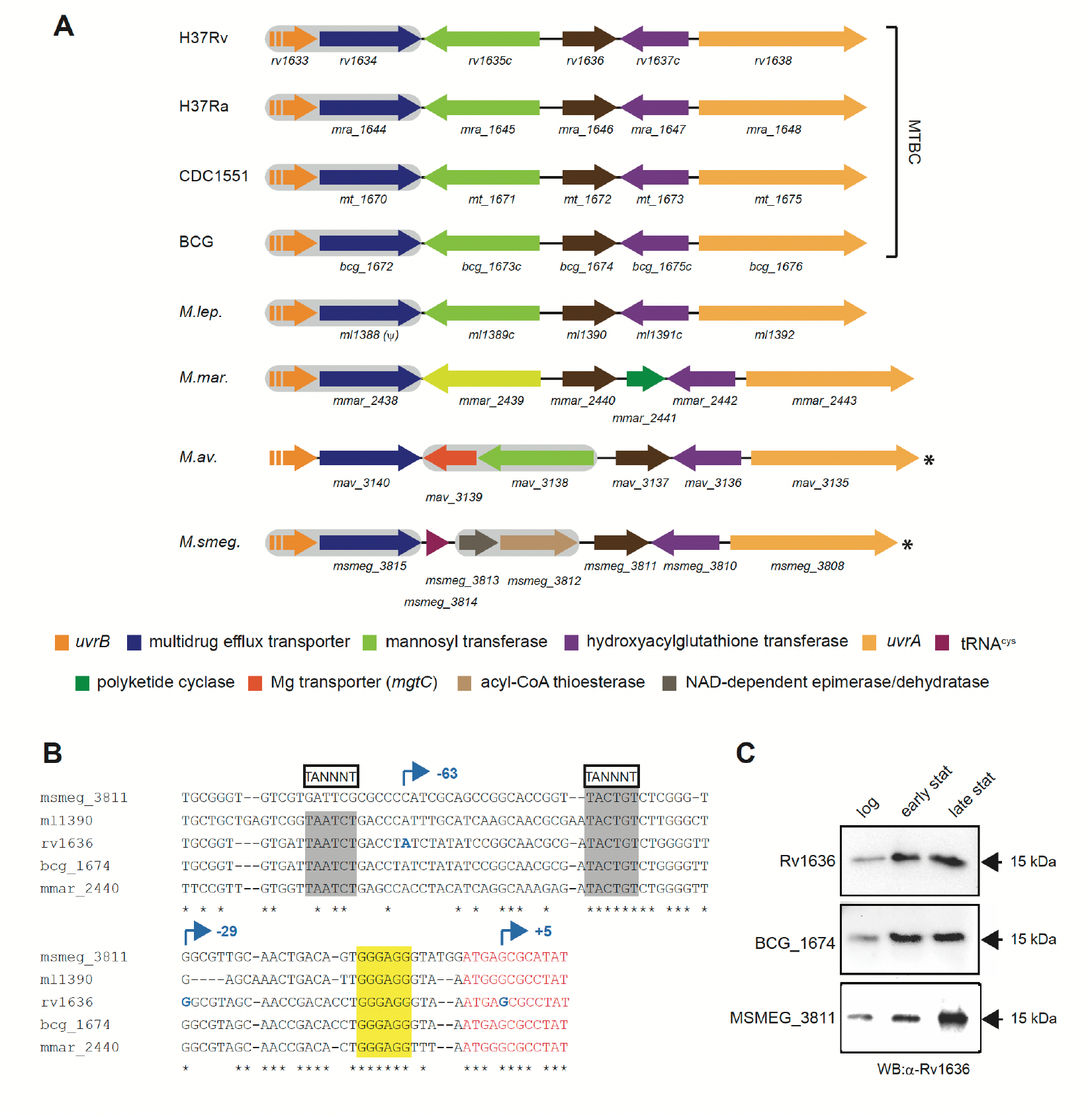
Genomic organisation, promoter architecture, and expression of *rv1636* and its orthologs. (A) Schematic overview of the locus encoding *rv1636* and its mycobacterial orthologs. Orthologs are similarly colour-coded with arrowheads indicating gene orientations. Genes predicted to occur in operons are highlighted in grey. A discontinuous arrow indicates partial representation of the gene. ψ indicates pseudogene, asterisks indicate genomic orientation in the negative strand. Gene lengths are not to scale. MTBC, *M. tuberculosis* complex; H37Rv, *M. tuberculosis* H37Rv; H37Ra, *M. tuberculosis* H37Ra; CDC1551, *M. tuberculosis* CDC1551; BCG, *M. bovis* BCG; *M.leps., M. leprae* TN; *M.mar., M. marinum; M.av., M. avium; M.smeg., M. smegmatis* mc^2^ 155. (B) Multiple sequence alignment of the promoters of *rv1636* and its orthologs. 85 nucleotides (black) upstream to the ORFs and 12 nucleotides (red) from the start of the ORFs have been taken for the alignment. The mapped TSSs are indicated by arrows (blue) and their respective positions with respect to the start codon of *rv1636* are shown. The putative −10 elements (TANNNT) and the putative Shine-Dalgarno sequence are highlighted in grey and yellow, respectively. (C) Expression of Rv1636, BCG_1674 and MSMEG_3811 as a function of growth. Immunoblot performed using antisera raised against recombinant Rv1636. Log, logarithmic; stat, stationary. Data shown is representative of experiments performed thrice. **Figure 1 Source Data 1. Source Data 1C, western blots shown in Figure 1C.**

### Expression of Rv1636/MSMEG_3811 during mycobacterial growth

Based on two earlier reports of transcription start site (TSS) mapping from *M. tuberculosis* using RNA-sequencing (31, 32), three possible TSSs for *rv1636* were identified, located at - 63, −29 and +5 with respect to the translation start codon, respectively (Fig 1B). The −63 site was ascribed as the primary TSS based on the peak intensities (31, 32). Interestingly, the transcript originating at +5 position would give rise to a leaderless variant, whose translated product would lack the first 6 amino acids in comparison to the full-length protein translated from the other two leadered transcripts. In addition to the TSSs, the −10 element consensus identified in mycobacteria - ‘TANNNT’ (31) - was found to be conserved for both the −63 and −29 TSSs across diverse mycobacterial species, except for *msmeg_3811* where the TSS at −29 appeared to be the primary one (Fig 1B).

We monitored the expression of Rv1636 and its orthologs from two mycobacterial species, *viz. M. bovis* BCG and *M. smegmatis*. Interestingly, the gene sequences of *rv1636* and *bcg_1674* and adjacent 1000 bp upstream and downstream are identical. Rv1636 levels increased as the cells entered stationary phase and the levels were maintained even in the late stationary phase (Fig 1C). This pattern of expression is common across USPs from diverse bacterial species (33, 34). BCG_1674 and MSMEG_3811 expression also followed a similar pattern (Fig 1C).

### Secretion of Rv1636 is SecA2-dependent

Prior to our identification of Rv1636 as a cAMP-binding protein, several independent proteomics-based studies had reported Rv1636 to be a secreted protein both *in vitro* and in animal models of infection, including sera from human smear-positive TB patients (35–40). Studies with *M. bovis* BCG also showed BCG_1674 to be secreted (36, 38). However, the mechanism of secretion of Rv1636 (and its orthologs) remained unclear.

Rv1636 lacks any canonical signal peptide at its N-terminus, ruling out secretion via the SecA1 and Tat pathways. Rv1636 is not located in any of the ESX-encoding loci in addition to being functionally unlinked to most of the ESX pathways. Further, the fact that *M. bovis* BCG, a strain inherently lacking the ESX-1 system (RD1 locus) also secretes Rv1636 (BCG_1674), rules out the involvement of ESX-1. Reported substrates of the SecA2 pathway in *M. tuberculosis, viz*. SodA and KatG, are known to lack a canonical signal peptide (41, 42). We therefore asked whether Rv1636 secretion was SecA2-dependent.

We probed the culture filtrate fractions from *M. tuberculosis* wildtype, *ΔsecA2* and *ΔRD1* strains, respectively, by immunoblotting using anti-Rv1636 antisera (Fig 2A). Rv1636 could be readily identified in the culture filtrate fractions of wildtype and *ΔRD1* cells, but was absent from the culture filtrate fraction of *ΔsecA2* strain (Fig 2A). CRP antibodies were used to confirm that the protein seen in the culture filtrate was not a consequence of cell lysis, since CRP is exclusively an intracellular protein (Fig 2A). We noted that secreted Rv1636 migrated faster than its cytosolic counterpart (Fig 2B). N-terminal sequencing of secreted Rv1636 had demonstrated the absence of the first 6 amino acids (36), but this may not lead to a significant change in size of the protein in culture supernatants, as is seen in blots. Moreover, no evidence of the shorter form was seen in the cytosol. Therefore, it is possible that secretion-dependent protein processing occurs.

**Figure 2.**
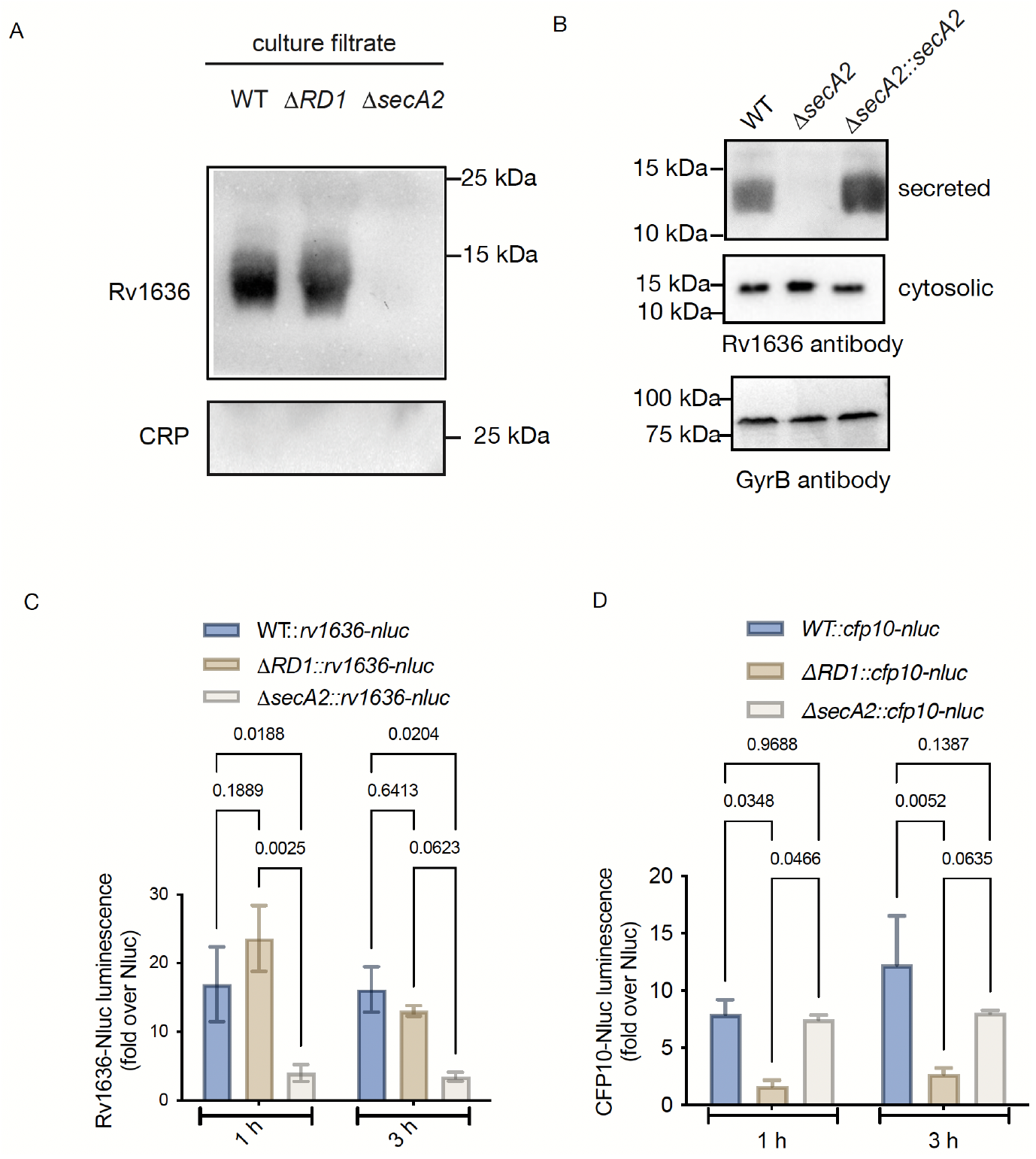
Secretion of Rv1636 is SecA2-dependent. (A) Immunoblots probing for Rv1636 and CRP in the culture filtrate fractions prepared from *M. tuberculosis* wildtype, *ΔRD1* and *ΔsecA2* strains. (B) Rv1636 levels were measured in the culture filtrate (secreted) and cytosol as detected by immunoblotting in *M. tuberculosis* wildtype, *ΔsecA2* and *ΔsecA2::secA2* complemented strains, respectively. Cytosolic protein loading was normalized to gyrase B levels. Data shown in A-B are representative blots of experiments performed thrice. (C-D) Secretion of CFP-10 and Rv1636 monitored using *cfp10-nluc* and *rv1636-nluc* fusions, respectively, where the corresponding fusion constructs were expressed under the *cfp-10* and *rv1636* native promoters. Relative CFP-10 and Rv1636 secretion across the three strains expressed after normalisation with Nluc bioluminescence in the culture filtrates. Mean ± SD plotted for duplicate cultures across two biological replicates. Two-way ANOVA with Bonferroni post-test and p values are shown. Data is representative of experiments repeated twice with duplicate determinations at each time point. **Figure 2 Source Data. Source Data 2A, western blots shown in Figure 2A. Source Data 2B, western blots shown in Figure 2B.**

Complementation of SecA2 in the *ΔsecA2* strain restored Rv1636 expression in the culture filtrate confirming that secretion of Rv1636 was mediated by SecA2. We also observed that the cytosolic levels of Rv1636 appeared to be higher in the *ΔsecA2* strain compared to both wildtype and *ΔRD1* (Fig 2B), suggesting intracellular accumulation of the cytosolic variant in the absence of secretion.

We developed a novel nanoluciferase (NanoLuc/Nluc)-based assay to monitor protein secretion (22). First, to validate this assay, we generated a fusion construct of CFP-10 and nanoluciferase (CFP10-Nluc) using a flexible linker (22), and introduced it into both wildtype, *ΔRD1* and *ΔsecA2* strains. Secretion of CFP-10 is dependent on the *RD-1* locus. We monitored the nanoluciferase activity in the culture supernatants from these strains and found a significant reduction in luciferase activity in the culture supernatant from the *ΔRD1* strain (Fig 2C), establishing that our assay could effectively reflect protein secretion in mycobacterial cells. Next, we tested secretion of Rv1636 using a Rv1636-Nluc construct. At both 1 and 3 h timepoints, nanoluciferase activity appeared to be significantly lower in the culture supernatant of the *ΔsecA2* strain compared to both wildtype and *ΔRD1* (Fig 2D). Therefore, we conclude that secretion of Rv1636 is SecA2-dependent.

### Levels of Rv1636/MSMEG_3811 and intracellular cAMP

We next quantified the amount of intracellular MSMEG_3811 in wildtype *M. smegmatis* cells using known amounts of recombinant MSMEG_3811 and densitometric analysis of bands seen on Western blotting (Fig 3A). In logarithmic and stationary phases of growth, the absolute levels of MSMEG_3811 was estimated as ~1 and ~3 pmoles per 100 μg of total protein, respectively (Fig 3A). However, the total and bound cAMP levels in *M. smegmatis* were significantly higher than levels of MSMEG_3811 (Fig 3A, lower panel), suggesting that additional cAMP-binding proteins could contribute to the bound cAMP fraction.

**Figure 3.**
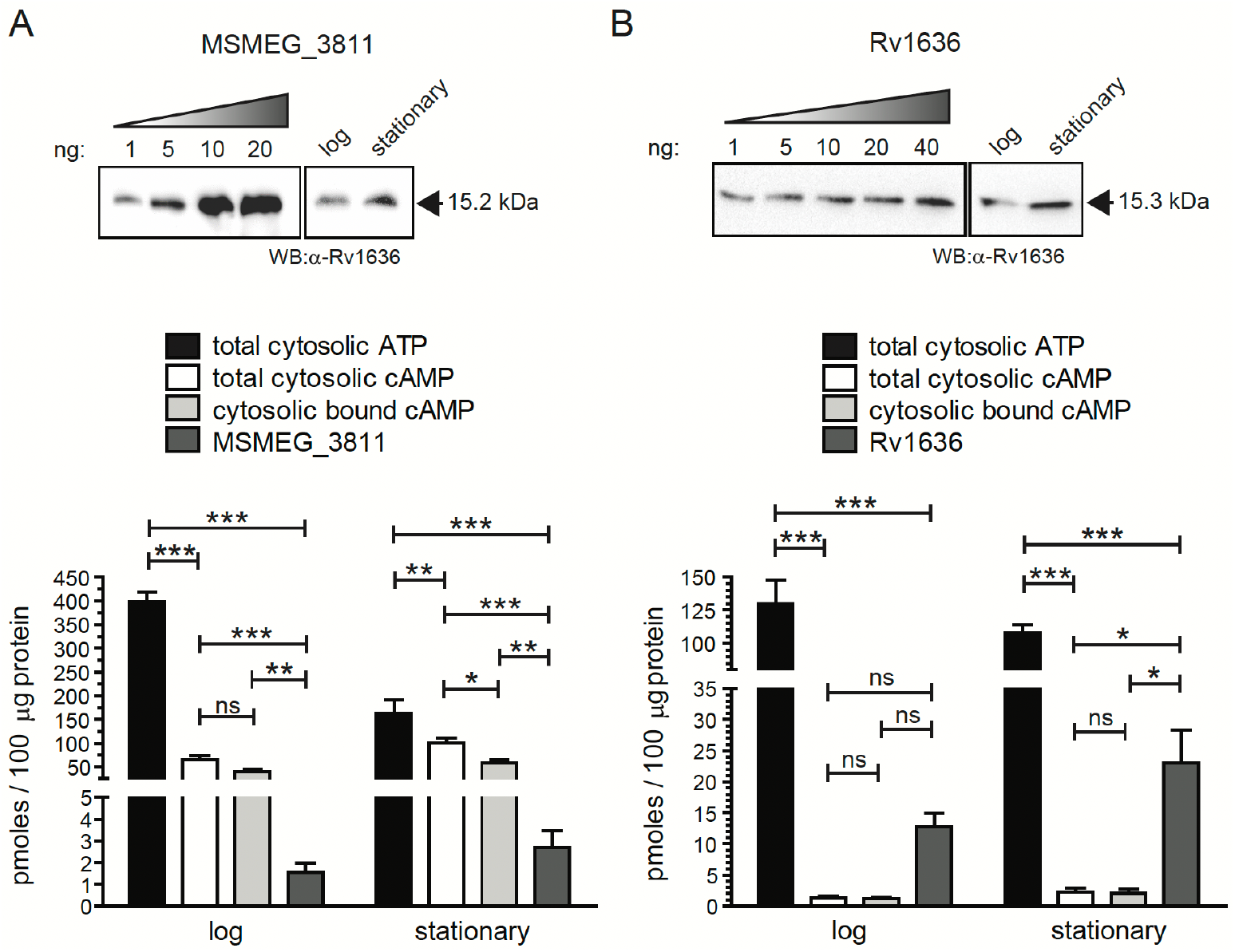
Determination of cytosolic levels of Rv1636, MSMEG_3811 and intracellular cAMP. (A-B) Top panels show intracellular levels of MSMEG_3811 and Rv1636 in *M. smegmatis* and *M. tuberculosis*, respectively, estimated by interpolating from standard curves generated by using known amounts of recombinant MSMEG_3811 (A) or Rv1636 (B) and densitometric analysis following immunoblot with anti-Rv1636 antisera. Cells from both logarithmic and stationary phases of growth were taken, and data shown is representative of experiments performed twice. Bottom panels show amounts of Rv1635/MSMEG_3811, amounts of cAMP (total and bound) and ATP (total) levels for direct comparison. Intracellular protein, total and bound cAMP, and total ATP were measured from the same lysates with duplicate determinations. One-way ANOVA with Tukey’s multiple comparison post-test performed for log and stationary phases, respectively; ns represents p values >0.05, * represents p value <0.05, ** represents p value <0.01, and *** represents p valuse <0.001. Data shown is from experiments performed at least twice in duplicate. **Figure 3 Source Data. Source Data 3A, western blots shown in Figure 3A. Source Data 3B, western blots shown in Figure 3B.**

In slow-growing pathogenic *M. tuberculosis*, levels of intracellular Rv1636 ranged from ~10-20 pmoles per 100 μg of total protein between logarithmic and stationary phases of growth (Fig 3B). Interestingly, cAMP levels in *M. tuberculosis* appeared to be much lower, ranging between ~1-2 pmoles per 100 μg of protein (Fig 3B). In contrast, levels of ATP, a lower affinity ligand for Rv1636 (10), remained significantly higher than Rv1636 and cAMP (Fig 3A-B). Similar to findings in *M. bovis* BCG (10), almost the entire fraction of intracellular cAMP in *M. tuberculosis* was protein bound (Fig 3B). suggesting that Rv1636 could be the major cAMP-binding protein in this species. Consequently, the amount of free cAMP available to other cAMP effectors in slow-growing mycobacteria, including *M. tuberculosis*, may be tightly regulated by levels of Rv1636 in the cell.

### Rv1636/MSMEG_3811: an intracellular ‘sink’ for cAMP

Cyclic AMP-binding to the GAF or CNB domains in proteins relays the signal to an associated effector domain that carries out downstream functions in a signaling cascade (12–14). In contrast, in Rv1636/MSMEG_3811, the USP domain to which cAMP binds is not associated with any other functional or catalytic domain. We therefore hypothesised that Rv1636/MSMEG_3811 may act to sequester the cyclic nucleotide. In contrast to slow-growing *M. bovis* BCG (10) and *M. tuberculosis* (Fig 3B), where almost the entire cytosolic cAMP was protein bound, fast-growing *M. smegmatis* could be used to test our hypothesis, since only ~50 % of the intracellular cAMP in this species exists as a bound fraction (Fig 3A).

Overexpression (OE) of Rv1636 under its own promoter from a low-copy number episomal plasmid (Fig 4A) revealed no difference in the growth of the cells with respect to the vector control (Fig 4B). We separated the intracellular cAMP into protein bound and free fractions (Fig 4C). The total intracellular cAMP levels between the Rv1636 overexpressing strains and the vector control remained comparable, however, the bound cAMP levels approximately doubled in the OE strain (Fig 4C). This significant increase in the bound cAMP levels represented an increase of ~40-45 pmoles of bound cAMP per 100 μg of protein (Fig 4C). Interestingly, the total extracellular cAMP levels remained unchanged (Fig 4D).

Conversely, we argued that deletion of *msmeg_3811* gene would result in a loss or decrease of bound cAMP levels in *M. smegmatis* cells. Therefore, we generated an unmarked deletion strain of *msmeg_3811* (*Δmsmeg_3811*) (Fig 5 A-B). Deletion of *msmeg_3811* did not have any effect on either the growth (Fig 5C) or cell viability (Fig 5D) of *M. smegmatis* under the conditions of growth used in our experiments. However, the intracellular bound cAMP levels in *Δmsmeg_3811* were found to be marginally lower than the wildtype cells despite the total cAMP levels being similar (Fig 5E). This ~20 % decrease in the intracellular bound cAMP levels, represented ~10 pmoles of cAMP per 100 μg of protein (Fig 5E). This increased abundance of free cAMP in the *Δmsmeg_3811* cells was reflected by a marginal increase in total extracellular cAMP levels (Fig 5F). Taken together, overexpression of Rv1636 and deletion of *msmeg_3811* provide support for our hypothesis that Rv1636/MSMEG_3811 may sequester cAMP thus regulating or tuning free cAMP concentrations in the cells.

**Figure 4.**
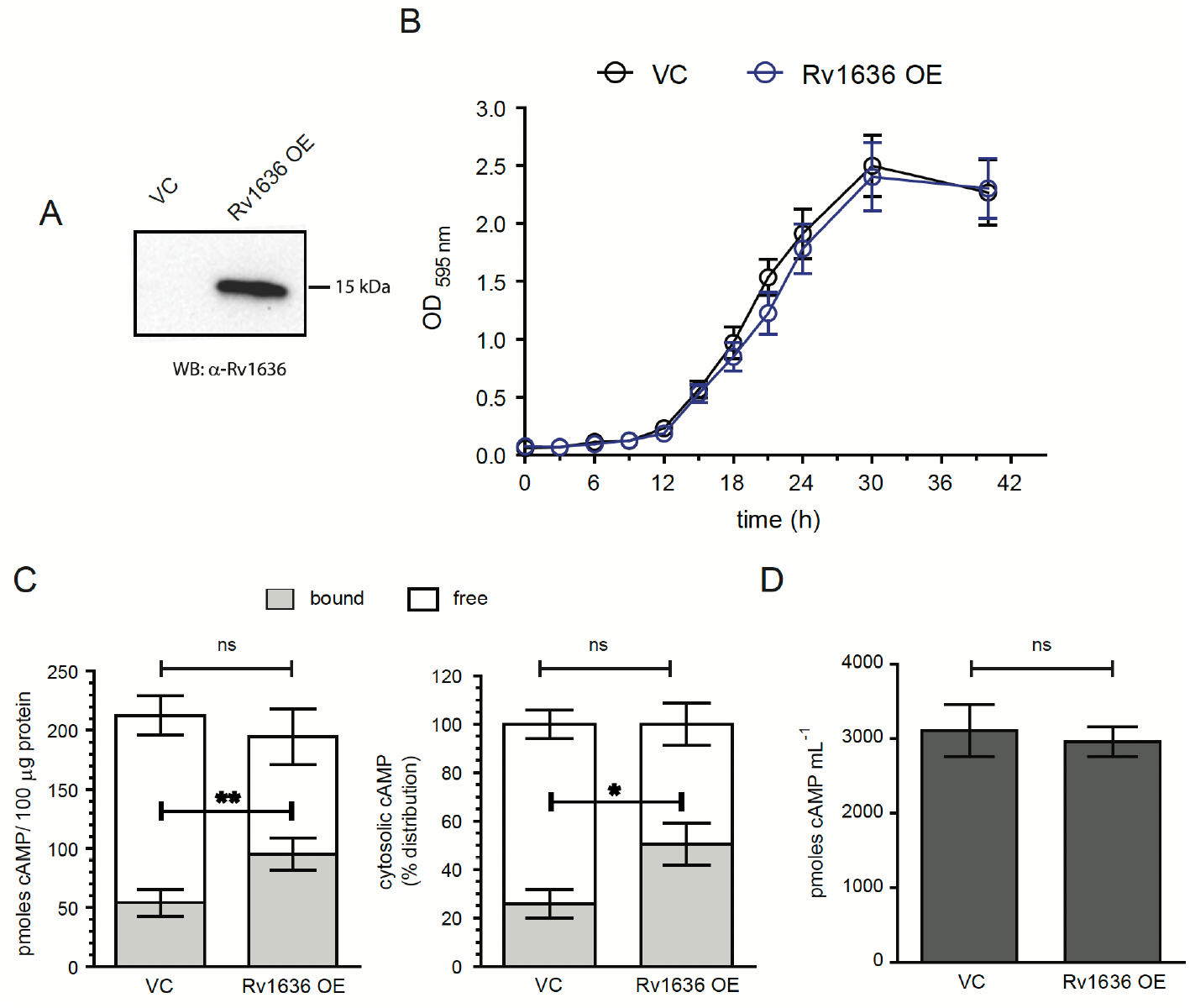
Overexpression of Rv1636 increases bound intracellular cAMP levels in *M. smegmatis*. (A) Immunoblot showing overexpression of Rv1636 in wildtype *M. smegmatis* mc^2^ 155. Blot shows lysates prepared from *M. smegmatis* cells transformed with either a vector control (VC) or Rv1636 overexpressing plasmid (OE). The blot shown is representative of two independently grown colonies. (B) Growth curve of Rv1636 OE (blue) versus VC (black). Cultures in duplicate were grown for the indicated time, and aliquots taken for measuring OD at 595 nm. Mean ± SD is plotted for experiments performed twice. (C) Plot showing the distribution of bound and free cAMP in the intracellular fractions of Rv1636 OE and VC strains. The sum of the free (white) and bound (grey) cAMP levels represents the total cytosolic cAMP levels. Left-hand panel shows the distribution of the free and bound cAMP in the intracellular fraction expressed as pmoles of cAMP per 100 μg of protein. Right-hand panel shows the same distribution of total intracellular cAMP into free and bound fractions as a percentage of the total. Cultures were grown in duplicate and data for experiments performed twice is shown. Paired t-test was performed for bound and free cAMP/% distribution independently; * represents p value <0.05, and ** represents p value <0.01. (D) Extracellular cAMP levels measured from the culture supernatants of Rv1636 OE and VC strains. Data represent mean ± SEM from two biological replicates, each having a pair of technical duplicates. Paired t test; p value >0.05 considered as non-signifìcant (ns). **Figure 4 Source Data. Source data 4A, western blot shown in Figure 4A.**

**Figure 5.**
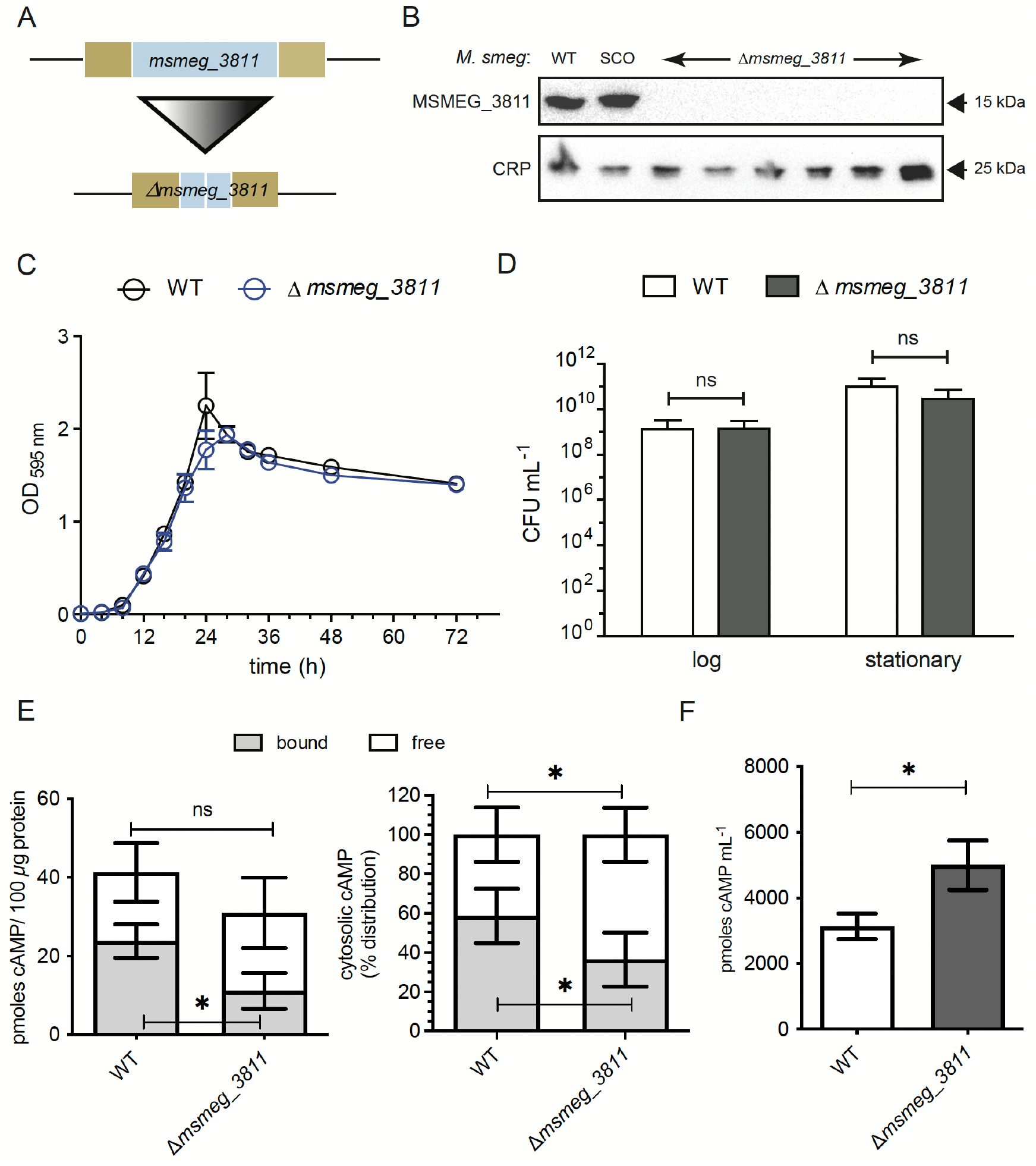
Deletion of *msmeg_3811* leads to reduced bound intracellular cAMP levels. (A) Schematic depiction of the wildtype *msmeg_3811* (top) and *Δmsmeg_3811* (bottom) alleles. The blue bar represents the *msmeg_3811* ORF in both the wildtype and *Δmsmeg_3811* strains. The brown bars represent the flanking sequences used for the homologous recombination. Image not to scale. (B) Immunoblot confirming the *Δmsmeg_3811* strains by probing with anti-Rv1636 antisera. Wildtype and SCO strains used as a positive control. Probing with anti-CRP antibody used as a positive control for all *M. smegmatis* strains. SCO, single crossover. Data shown is representative of blots performed twice. (C) Growth analysis of wildtype (WT) and *Δmsmeg_3811* strains. Mean ± SEM for two biological duplicates plotted from a representative experiment performed twice (D) CFU enumeration from logarithmic and stationary phase cultures of wildtype and *Δmsmeg_3811* strains. Mean ± SD plotted of a representative experiment performed twice, with cultures grown in duplicate. CFU numbers were analysed by independent t-tests with log and stationary cultures. (E) Bound and free cAMP levels in the intracellular fractions of wildtype and *Δmsmeg_3811* strains. Left hand panel shows levels of cAMP, right hand panel shows percentage distribution of bound and free cAMP of the total intracellular levels. Data shown is the mean ± SD from two independent experiments performed with duplicate cultures. (F) Extracellular cAMP levels in wildtype and *Δmsmeg_3811* strains. Mean ± SEM plotted from two biological duplicates for each genotype across multiple measurements. Paired t-test was performed; * represents p value <0.05. **Figure 5 Source Data. Source data 5B,Western blots shown in Figure 5B.**

### *rv1636* is an essential gene for survival of *M. tuberculosis*

We have seen that almost all cAMP in slow growing mycobacteria remained bound to protein, and concentrations of Rv1636 were high enough to serve as the cAMP sink in these bacteria (Fig. 3B). While the deletion of MSMEG_3811 would have only marginally increased free cAMP in the cell, the consequences resulting from deletion of *rv1636* may be more profound, if Rv1636 was responsible for maintaining ‘bound’ levels of cAMP. We therefore attempted to generate an unmarked deletion of *rv1636* in *M. tuberculosis* using a two-step, homologous recombination-based strategy (Fig 6A) (26). We confirmed two independent single crossover (SCO) colonies for the *rv1636* locus in wildtype *M. tuberculosis* by both PCR and Southern blotting (Fig 6A, right panel). Once the SCOs were confirmed, they were grown for the second crossover to occur followed by subsequent selection of the second crossovers (26). Since the two crossover events are selected in a stepwise manner independent of each other, the expected outcome after the second crossover is to get both wildtype and knockout colonies in a 1:1 ratio (26). Surprisingly, all the 47 legitimate double crossover (DCO) colonies screened appeared to be wildtype, suggesting that *rv1636* may be an essential gene.

We generated a merodiploid strain (*Mtb::rv1636*) where we introduced a second functional, wildtype copy of *rv1636* at the mycobacteriophage L5 *att* site under its own promoter (Fig 6A). Rv1636 levels were detected by immunoblotting (Fig 6B). We then deleted the copy of *rv1636* at the endogenous locus (Fig 6A) in the merodiploid strain. We were readily able to generate a deletion of *rv1636* at the endogenous site (Fig 6C), while the L5 *att* site copy remained intact (Fig 6C).

In another approach, we designed a CRISPRi-mediated knock down of *rv1636* in wildtype *M. tuberculosis* cells (27). The sgRNA specific to *rv1636* could anneal from the +11 position in the ORF, halting gene transcription. When the *rv1636*-specific sgRNA was induced with anhydrotetracycline, the cells showed a severe growth retardation as opposed to scrambled sgRNA (Fig 6D). We monitored expression of Rv1636 by western blotting with lysates harvested on days 2 and 4 after anhydrotetracycline addition and observed almost complete depletion of Rv1636 within 48h. Together, these results indicate that *rv1636* gene was essential for the survival of *M. tuberculosis*.

Next, we asked whether the essentiality of *rv1636* was linked to its ability to bind cAMP. The crystal structure of the cAMP-bound MSMEG_3811 (PDB ID 5AHW) aided in the identification of residues that made direct contacts with the ligand (10). Gly-10 and Gly-114 make interactions with the 2’-OH of the ribose moiety and the cyclic phosphodiester, respectively (Fig 6F inset), and both the corresponding single mutants in Rv1636 - G10T and G113A - showed reduced cAMP binding (10). For this study, we mutated both these residues to generate a double mutant (Rv1636^DM^) variant. The recombinant Rv1636^DM^ protein lost all cAMP binding, as monitored using biolayer interferometry (BLI) (Fig 6F).

We generated a merodiploid strain (*Mtb::rv1636DM*) where we provided the *rv1636DM* allele as a second copy at the mycobacteriophage L5 *att* site. The merodiploid strain was confirmed by PCR and Southern blotting (Fig 6A) and the expression of Rv1636DM was quantitated by immunoblotting (Fig 6B). We tested whether deletion of the endogenous *rv1636* allele was possible in the background of the *rv1636DM*. Of a total of 23 legitimate DCO colonies screened, no knock-out was obtained. Therefore, this genetic evidence allows us to conclude that the essentiality of *rv1636* was dependent on its ability to bind cAMP.

### A natural compound identified by molecular docking inhibits *M. tuberculosis* growth

We have earlier described the crystal structure of MSMEG_3811 bound to cAMP, and structure-guided mutational analysis identified residues critical for cAMP binding (10).

Further, using computational docking and library screening approaches, of the two natural compounds that could bind to MSMEG_3811(43), STOCKIN42384 (Figure 7A) had a docking score better than that of cAMP, and engaged Ala40 (a key residue for cAMP binding (10)) and Glu57 in hydrogen bonding. The second compound, STOCKIN74667 (Figure 7A), was hydrogen bonded to Gly114, also important for cAMP binding. Both these compounds were shown to bind to Rv1636 with K_d_ ~ 1 mM (43). We tested the ability of *M. tuberculosis* to grow in the presence of both these compounds.

**Figure 6.**
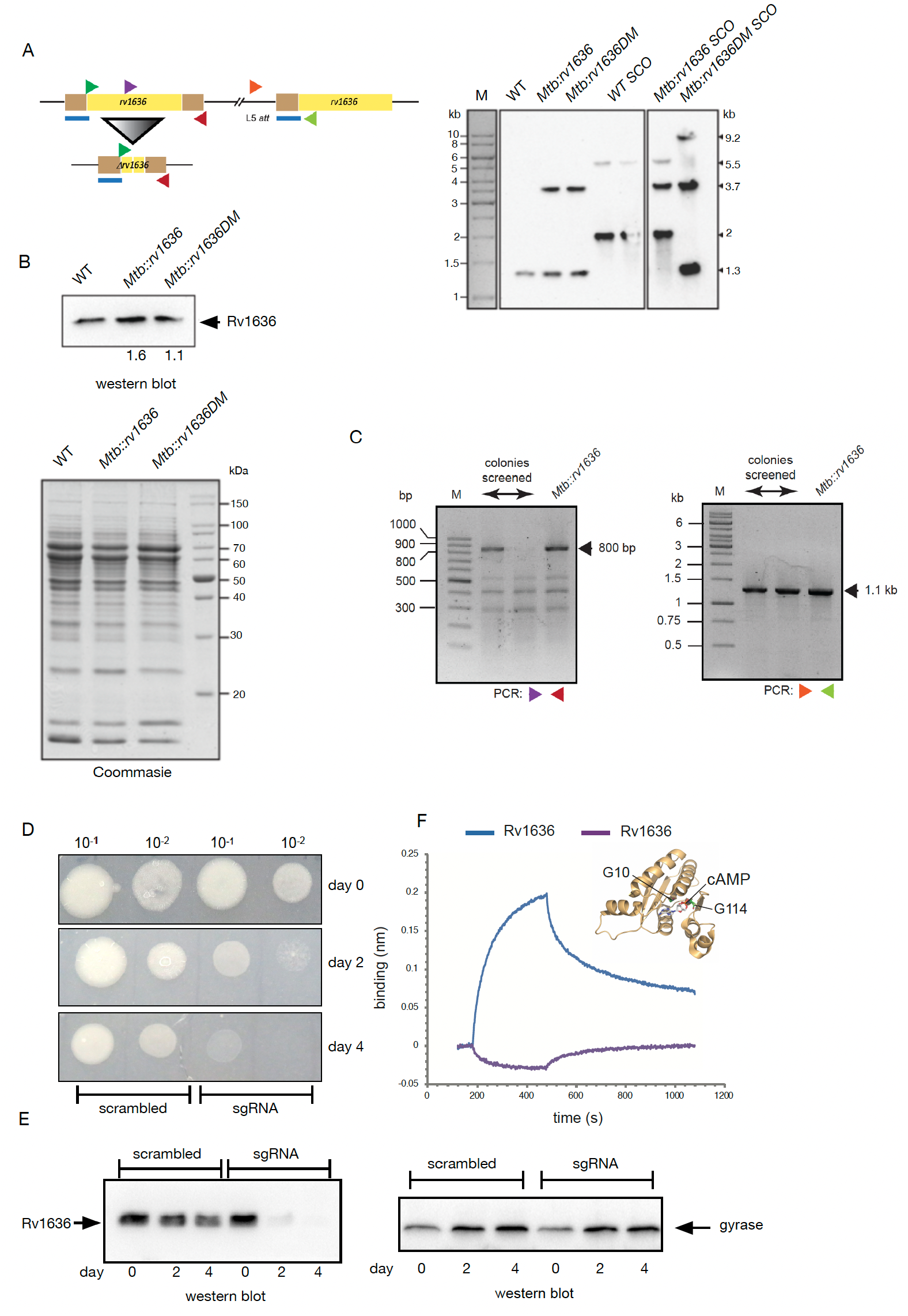
The *rv1636* gene is essential in *M. tuberculosis* and essentiality is dependent on its cAMP-binding ability. (A) Schematic representation of the wildtype and *Δrv1636* alleles, the yellow bar representing the gene ORF while the flanking genomic regions are depicted in brown. At the L5 *att* site, the additional *rv1636* allele containing ~650 nt upstream region (brown) along with the ORF (yellow) represents the DNA fragment used to construct the merodiploid strains. The blue line indicates the region used to probe the *rv1636* allele in Southern blot. The PCR primer pairs used in screening (Fig 6C) are depicted as colour-coded arrowheads. Image not to scale. Right panel: Southern blot to confirm the genotypes of the following *M. tuberculosis* strains – wildtype (WT), merodiploid having an additional copy of either the wildtype (*Mtb::rv1636*) or double mutant *rv1636* (*Mtb::rv1636DM* alleles, wildtype single crossovers (WT SCO), and single crossovers in the merodiploid backgrounds, respectively. The number of alleles in the respective strains are detected by the number of bands at their corresponding sizes. Genomic DNA was digested with *Kpn*I prior to transfer and blotting. (B) Immunoblot showing the levels of expression of Rv1636 in the merodiploid strains *Mtb::rv1636* and *Mtb::rv1636^DM^*, respectively, compared to wildtype *M. tuberculosis*. Numbers below the blot represent the relative densitometric amounts across two experiments. Bottom panel shows SDS-PAGE profiles of the lysates used for immunoblotting from a representative experiment. (C) PCR to confirm DCO colonies from *Mtb::rv1636* where only the wildtype allele at the endogenous locus will generate a 800 bp product. Absence of this product confirms the deletion of the endogenous *rv1636* gene (third lane from the left). Right panel: PCR to confirm the presence of the *rv1636* allele at the L5 *att* site in the DCO colonies shown in Fig 6C left panel. (D) CRISPRi (*Spy* dCas9)-mediated knock-down of *rv1636* using a *rv1636*-targeting sgRNA which binds spanning the +11 to +30 position within the *rv1636* ORF. Upon induction of CRISPRi in the presence of Atc, severe growth retardation was observed only with the specific sgRNA targeting *rv1636* (bottom right). Data shown is representative of experiments performed thrice. (E) Immunoblot showing depletion of Rv1636 protein levels as a function of Atc treatment duration with *rv1636*-specific sgRNA as opposed to scrambled control. Blot is representative of lysates from a single experiment, with the experimental knockdown performed thrice. (F) Cyclic AMP binding of Rv1636 and Rv1636^G10T G113A^ (Rv1636^DM^) monitored using bio-layer interferometry. Inset showing the crystal structure of a cAMP-bound MSMEG_3811 monomer (from PDB ID 5AHW), indicating the positioning of G10 and G114 (equivalent of G113 in Rv1636) residues in the cAMP-binding pocket. Data shown is representative of experiments performed twice with individual protein preparations. **Figure 6 Source Data 6. Source Data 6A, Southern blots shown in Figure 6A. Source Data 6B, Coomassie stained gel and western blots shown in Figure 6B. Source Data 6C, PCR gels for data shown in Figure 6C. Source Data 6E, western blots shown in Figure 6E.**

**Figure 7.**
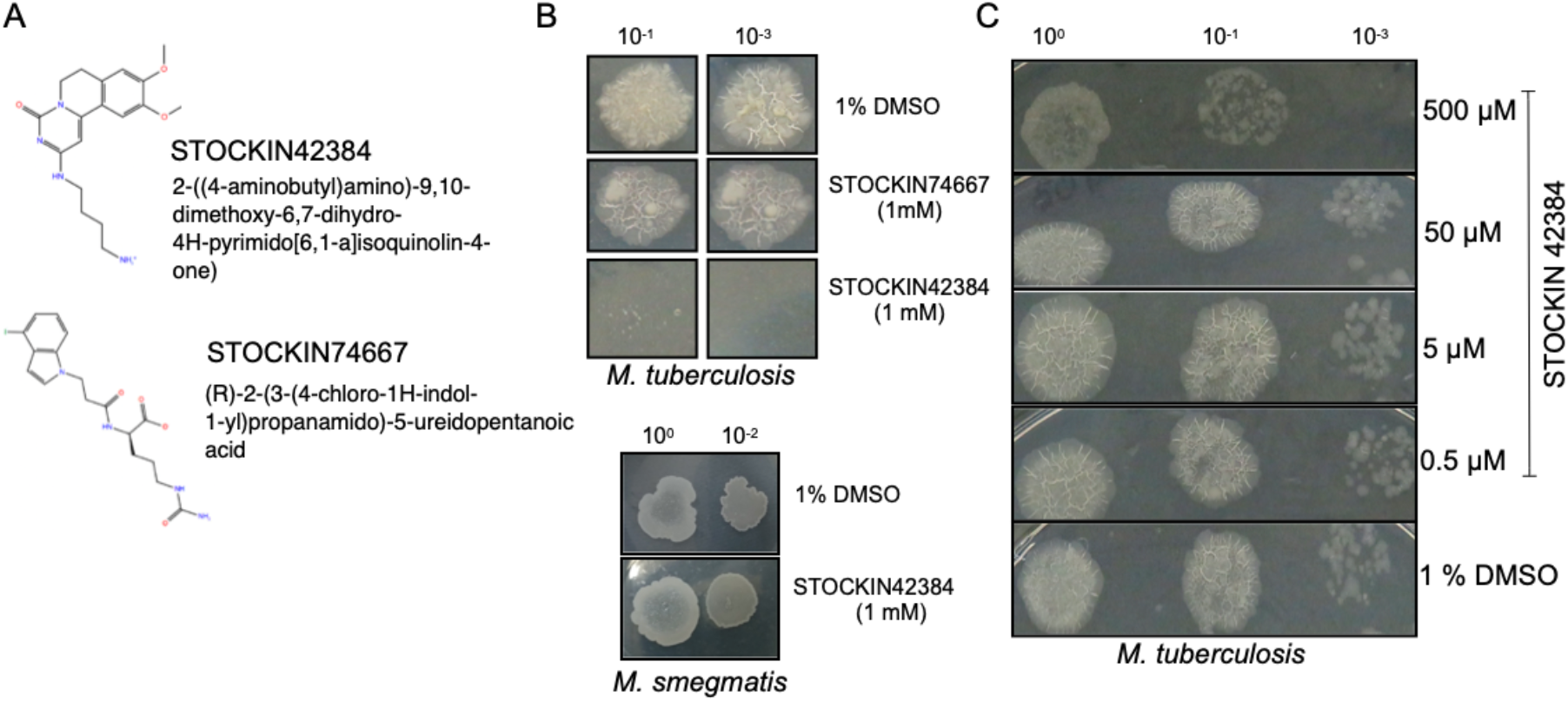
Inhibition of growth of *M. tuberculosis* by a natural compound that binds Rv1636. (A) Structure of the two compounds identified by molecular docking (43). (B) Growth of *M. tuberculosis* and *M. smegmatis* in media containing 1mM of the two natural compounds, or 1% DMSO that was used as the solvent to dissolve the compounds. Image shown is from an experiment from two independent cultures spotted in duplicate (C) Growth of *M. tuberculosis* in the presence of indicated concentrations of STOCKIN42384. The experiment was repeated twice.

As shown in Figure 7B, complete inhibition of growth of *M. tuberculosis* was seen when grown in the presence of STOCKIN42384 while growth was similar in media containing STOCKIN74667 and media containing DMSO (solvent used to dissolve the compounds) alone. Importantly, there was no growth inhibition of *M. smegmatis* with STOCKIN42384. This suggests that STOCKIN42384 binding to Rv1636 (with a docking score of −11.1 kcal/mole versus −10.5 for cAMP) prevents cAMP binding and mimics the knock-down of this protein encoded by an essential gene in *M. tuberculosis* (Figure 6). The non-essentiality of MSMEG_3811 allows the growth of *M.smegmatis* in the presence of STOCKIN42384 and indicates that the killing seen for *M. tuberculosis* was not mediated by a general toxicity of this compound to mycobacteria. Importantly, this data provides evidence that inhibitors of Rv1636 may be worthy of developing in future as a therapeutic strategy to combat tuberculosis.

## DISCUSSION

Mycobacterial genomes encode a complex cAMP synthesising as well as utilising machinery (3, 8), emphasizing the diversity in how this ancient small molecule is deployed in regulating diverse aspects of bacterial physiology. The addition of Rv1636/MSMEG_3811 to the existing group of cAMP-binding proteins is both unique and intriguing. It is a single-domain protein and binds cAMP via the USP-fold. Since the gene is encoded as a monocistron, its function may be unlinked from neighbouring genes at that locus. A total of 6 of the 10 USPs in *M. tuberculosis* fall under the DosR-regulon (44), and Rv1636 is not one of those, indicating its unique functional relevance.

The presence of the +5 TSS identifies *rv1636* as one of the 47 genes which can be transcribed as both a leadered and leaderless transcripts (31). The smaller secreted form of Rv1636 may be the translated product of the leaderless transcript or have undergone proteolytic cleavage during secretion. SodA, KatG, and components of the Mce transporter are known SecA2 substrates in *M. tuberculosis* (42, 45, 46) to which we now add Rv1636. In *M. marinum*, secretion of PknG is SecA2-dependent (47). The role of secreted Rv1636, however, remains to be elucidated. The fact that *secA2* mutant exhibits attenuated virulence (41, 42, 46, 48) warrants further investigation into the functions of secreted Rv1636.

We also developed a new assay to monitor protein secretion using nanoluciferase fusion constructs. Our assay could recapitulate findings by preparing proteins from the culture filtrate. In essence, this new nanoluciferase-based assay could be used in other bacterial systems as a readout for protein secretion both quantitatively and kinetically.

The absence of any associated domain together with the fact that Rv1636/MSMEG_3811 did not co-purify with any other major protein on cAMP-affinity chromatography (10), suggested that the role of Rv1636/MSMEG_3811 was to sequester cAMP in the cells. Both overexpression of Rv1636 and deletion of *msmeg_3811* in *M. smegmatis* provided support to this hypothesis. The mycobacterial replicon harboured in the episomal plasmid to overexpress Rv1636 typically has a copy number of ~5 in mycobacterial cells (49, 50). Given the similarities in biochemical properties of Rv1636 and MSMEG_3811 (10), an increase of ~40-45 pmoles of cAMP per 100 μg of protein in the bound fraction of cAMP on overexpression of Rv1636 correlates well with the reduction of ~10 pmoles of bound cAMP per 100 μg of protein in the *Δmsmeg_3811* cells.

The observation that neither overexpression of Rv1636 nor deletion of *msmeg_3811* had any effect on the growth of the *M. smegmatis* cells can be explained by the fact that the total, as well as free cAMP levels in fast-growing M. smegmatis, are far higher than MSMEG_3811 levels. In contrast, in slow-growing pathogenic *M. tuberculosis*, both intracellular cAMP and Rv1636 levels were comparable. An earlier report by Schubert *et al*. demonstrated that absolute levels of Rv1636 appeared to be 6.2 pmoles per 100 μg of protein, making it the 20^th^ most abundant protein in the cell (51). Interestingly, our genetic analyses showed that *rv1636* is essential for survival of *M. tuberculosis* and the ability of Rv1636 to bind cAMP was absolutely critical. This function is a new addition to the roles played by cAMP-binding proteins in regulating diverse physiological outcomes. It also suggests that Rv1636/MSMEG_3811 may function upstream of other canonical cAMP-binding proteins by limiting or releasing cAMP for binding to other cAMP-binding proteins.

Our finding that *rv1636* is an essential gene in *M. tuberculosis* is further supported by results from transposon insertion mutagenesis screens (52, 53). At least two such independent studies reported that of the 10 permissive TA insertion sites within the *rv1636* gene, only one insertion was observed after the TA of the stop codon in the ORF, essentially leaving the entire ORF unperturbed (52, 53). Interestingly, flanking genes of *rv1636* were not found to be essential. In the promoter and 5’-UTR of *rv1636*, TA sites are found at −3, −25, −43, −58, −60, −64 and −74 positions, respectively. Except for the −3 site, all sites had transposon insertions (52, 53). This could demarcate the minimal essential region for expression of *rv1636* which contained at least one transcription start site, which in this case was the +5 TSS. The study by Griffin *et al*. utilised minimal media plus glycerol (53), whereas DeJesus *et al*. had grown *M. tuberculosis* on 7H9 or 7H10 media containing oleic acid-albumin-dextrose-catalase (OADC) supplement (52), suggesting that the essentiality of *rv1636* may be independent of the growth media used or culture conditions.

None of the cAMP-binding protein encoding genes or other *usp* genes are predicted to be essential in *M. tuberculosis* (52, 53). The *crp/rv3676* gene is attenuated in its virulence (54), while the knockout of *kat/rv0998 in M. bovis* BCG shows a growth defect when grown in propionate as the sole carbon source (55). Individual deletions of 4 *usp* genes – *rv1996, rv2005c, rv2026c* and *rv2028c*, respectively, in *M. tuberculosis* H37Rv showed no growth disadvantage (56). Deletion of another *usp*, *rv2623*, in *M. tuberculosis* Erdman strain failed to establish chronic infection in guinea pig and mice models, showing a hypervirulence phenotype (57). This was linked to the ability of Rv2623 to be able to bind ATP (57). The ortholog of *rv1636* in *M. leprae, ml1390*, is the only functional *usp* gene encoded in its genome (56).

These observations indicate that Rv1636 may play a pivotal role in cAMP-mediated cellular signaling in Mycobacteria, by regulating levels of ‘free’ cAMP available for other effectors including CRP and the cAMP-dependent protein acyl transferase, Rv0998. This proposes a new mechanism of how cAMP-mediated signaling could be modulated in the cell, without changing absolute levels of cAMP in the cell (3, 8). High iron and treatment of ofloxacin and moxifloxacin have been reported to positively regulate Rv1636 levels (58, 59), and it would be interesting to measure ‘bound’ and ‘free’ levels of cAMP under these conditions. Interestingly, Rv1636 levels increase (Fig 1C) as mycobacterial cells start synthesising cAMP in the logarithmic phase of growth (2).

We have provided preliminary evidence that a compound identified by docking a natural library to the cAMP-bound structure of MSMEG_3811 could inhibit the growth of *M. tuberculosis* (Figure 7B, C). This compound (STOCKIN42384) *in vitro* showed low-affinity binding to Rv1636, but docking indicated a higher docking score than cAMP (43). Further, the fact that STOCKIN42384 bound in the cAMP-binding pocket of Rv1636 emphasizes that cAMP binding is an essential function of Rv1636 in *M. tuberculosis*. In our docking analysis, we could identify another compound, CHEMBL2109743 (GSK581005A) with a docking score of −10.9 kcal/mol, comparable to that for cAMP (−10.5 kcal/mol) (43). This compound has been shown to be effective against *M. bovis* BCG, with a ED50 of 9.8 μM (60), but the target to which it binds is not known. Testing this compound to check for binding to Rv1636 and effects on *M. tuberculosis* would be a worthwhile effort.

There are approximately 600-700 genes in *M. tuberculosis* genome which are predicted to be essential for survival of the bacteria, accounting for ~16 % of all genes (52, 53, 61). Inclusion of *rv1636* in this list suggests that Rv1636 may be a potent target for developing anti-mycobacterial drugs, given the complete absence of USPs in higher mammals. Our results with a natural compound that binds Rv1636 (Figure 7) strongly suggest that targeting Rv1636 may prove to be useful in designing novel therapeutics for the treatment of Tuberculosis.

## ACKNOWLEDGEMENTS

The authors acknowledge Dr. Ravi Manjithaya, Jawaharlal Nehru Centre for Advanced Scientific Research (JNCASR) India for kindly sharing plasmids and reagents used in this study); Dr. David Sherman, University of Washington, USA for the *ΔRD1* strain; Dr. William R. Jacobs, Albert Einstein College of Medicine, USA for the *ΔSecA2* strain; and Dr. Robert Hussion for the pRH2502 and pRH2521 plasmids.

A.B. is supported by Senior Research Fellowship from the Council of Scientific & Industrial Research, Government of India. S.S. is supported by Dr. D.S. Kothari Postdoctoral Fellowship, University Grants Commission, Government of India. A.S. is supported by a Wellcome Trust/DBT India Alliance Senior Fellowship (IA/S/16/2/502700). S.S.V. is a recipient of a J.C. Bose National Fellowship from the Department of Science & Technology, Government of India (SB/S2/JCB-18/2013) and is a Margdarshi Fellow supported by the Wellcome-Trust India Alliance (IA/M/16/1/502606). This work was supported by the Department of Biotechnology, Government of India (BT/PR15216/COE/34/02/2017) and funding from the DBT-IISc Partnership Program Phase-II (BT/PR27952/INF/22/212/2018/21.01.2019).

## MATERIALS AVAILABILITY STATEMENT

All newly derived strains and clones are available on request from the authors. Antibodies to Rv1636, gyrase and CRP are also available on request.

## DATA AVAILABILITY

All data generated or analysed during this study are included in the manuscript. Source data files have been provided for Figures 1–6.

